# Phase transitions in tumor growth IX: gastric cancer metastasis

**DOI:** 10.64898/2026.01.28.702339

**Authors:** J.M. Nieto-Villar, R. Mansilla

**Affiliations:** Department of Physical-Chemistry, A. Alzola Group of Thermodynamics of Complex Systems of M.V. Lomonosov Chair, Faculty of Chemistry, University of Havana, Cuba; Centro Peninsular en Humanidades y Ciencias Sociales, CEPHCIS, UNAM, México; A Centro de Investigaciones Interdisciplinarias en Ciencias y Humanidades, UNAM, México

**Keywords:** Gastric cancer metastasis, Phase transitions, Dissipation function, Nonlinear dynamics, Thermodynamic robustness, Dynamical systems, Chaotic attractors, Immune surveillance

## Abstract

We propose a novel three-compartment heuristic model that recasts gastric cancer metastasis into a framework of non-equilibrium thermodynamics and nonlinear dynamics. The system, encompassing primary, hepatic, and peritoneal tumor populations, exhibits a well-defined route to chaos: as immune surveillance weakens, the dynamics undergo a supercritical Andronov-Hopf bifurcation, giving rise to a limit cycle, followed by a Shilnikov-type saddle-foci bifurcation cascade leading to chaotic attractors. Our central finding is the introduction of a dissipation function, Ψ, constructed via a sensitivity-weighted, two-factor ansatz that integrates metabolic flux and dynamical influence. This spatially coarse-grained measure captures the system’s thermodynamic robustness. The analysis reveals a dynamical phase transition: while tumor aggressiveness peaks in the pre-metastatic limit-cycle regime, Ψ emerges as the definitive marker of the chaotic, treatment-resistant metastatic state, quantifying a sharp increase in systemic robustness that correlates decisively with advanced clinical stages (TNM III–IV). Consequently, this work provides a predictive framework grounded in the physics of metastasis, demonstrating that Ψ not only diagnoses but also defines the primary therapeutic target: the underlying thermodynamic robustness of the metastatic system. Thus, effective intervention must shift from merely reducing tumor mass to strategically destabilizing this robust dissipative structure, thereby preventing recurrence.

*PACS*: 05.45.-a; 87.18.-h; 87.19.xj; 05.70.Ln

**Highlights:**

- A novel three-compartment heuristic model reveals phase transitions and chaotic dynamics in gastric cancer metastasis.
- The dissipation function Ψ emerges as a quantitative thermodynamic metric of systemic robustness in the metastatic regime.
- Decreasing immune surveillance triggers biological phase transitions towards metastatic disease.
- The framework integrates nonlinear dynamics with TNM staging, identifying the dissipation function Ψ as a therapeutic target to overcome metastatic recurrence.

**Graphical Abstract:** 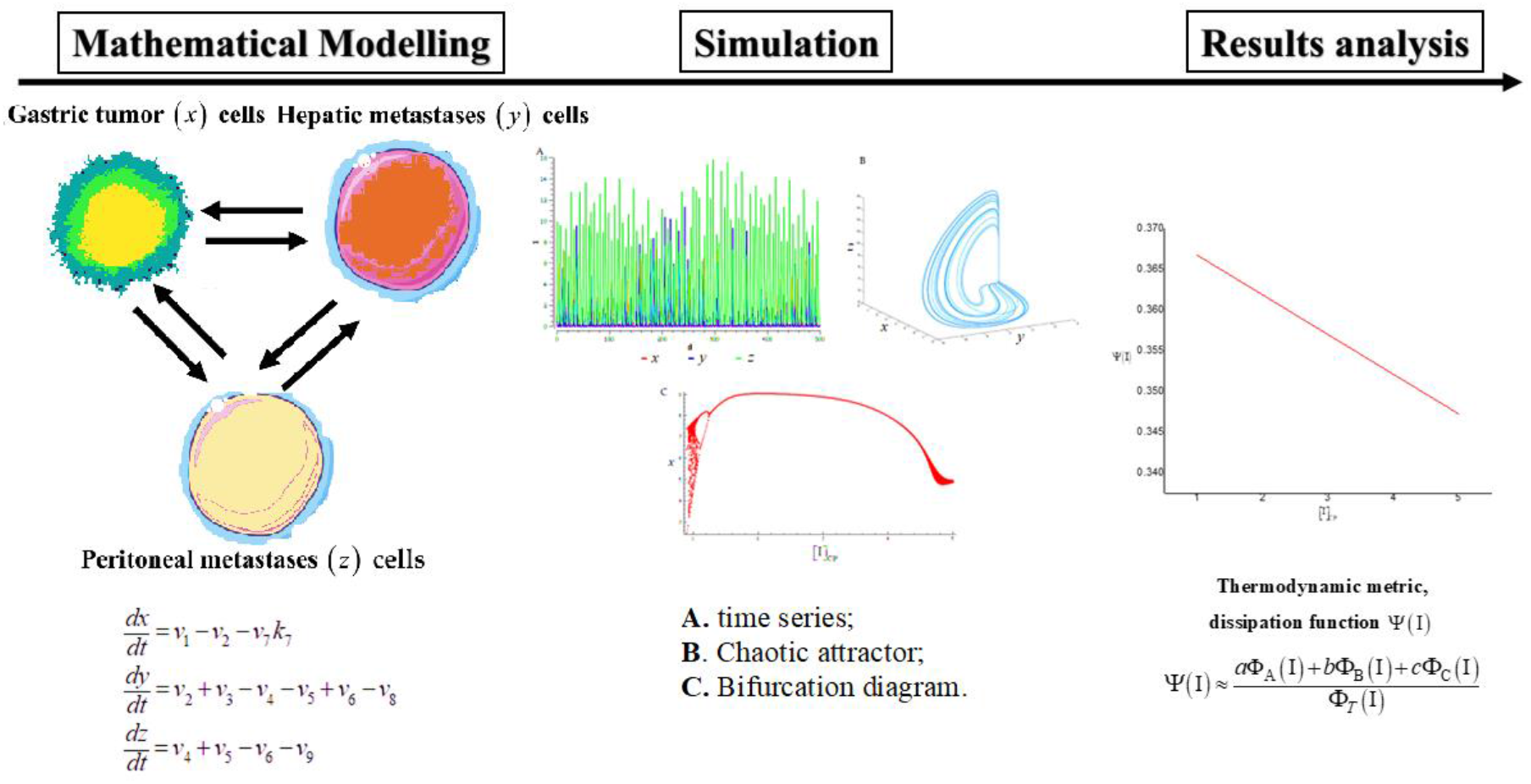

## 1. INTRODUCTION

Gastric cancer remains a major global health challenge, ranking as the fifth most common malignancy and the fourth leading cause of cancer-related death worldwide [1]. Its incidence shows significant geographic disparities, with the highest burden in Eastern Asia [2]. A central factor in its lethality is metastatic dissemination, particularly to the liver and peritoneum, which drastically reduces survival prospects and complicates therapeutic intervention [3,4]. Although a gradual decline in incidence has been attributed in part to factors like reduced *Helicobacter pylori* infection, the management of metastatic disease continues to represent a critical frontier in oncology [5].

Current efforts to predict and model gastric cancer metastasis predominantly employ statistical and machine-learning techniques [6,7]. These include logistic regression models, deep-learning algorithms trained on clinical imaging, and prognostic nomograms [8,9,10]. While valuable for risk stratification, these data-driven approaches are inherently descriptive and correlative. They lack a foundational theory that explains why metastasis occurs as a systemic, often sudden and irreversible, transition in the disease state. This creates a gap between prediction and a mechanistic, physics-based understanding.

An alternative paradigm posits that cancer is not merely an accumulation of mutated cells but a complex, self-organizing system operating far from thermodynamic equilibrium [11]. Substantial evidence supports this view [12,13], indicating that cancer emergence and evolution exhibit nonlinear dynamics, fractal structures, and deterministic chaos, which confer robustness, adaptability, and therapeutic resistance [14,15,16,17,18,19]. Within this framework, the progression from a localized tumor to systemic metastatic disease can be understood as a “biological phase transition”, a concept our group has advanced in previous work [20,21]. These transitions are driven by bifurcations in the underlying dynamical system, leading to new and robust organizational states [20,21].

Despite this theoretical progress, the application of non-equilibrium thermodynamics and nonlinear dynamics to the specific and clinically crucial problem of gastric cancer metastasis, particularly its dual routes to the liver and peritoneum, remains largely underdeveloped. It is paradoxical that, given the high burden of this disease, according to the best of our knowledge, how few works model its most lethal feature from a fundamental physics perspective [22,23,24,25,26].

Therefore, the objective of this work is to extend our established theoretical framework of phase transitions in tumor growth [11,27-34] to gastric cancer metastasis. We develop a novel three-compartment heuristic model to capture the nonlinear coupling between the populations of the primary gastric tumor (*x*), hepatic metastases (*y*), and peritoneal metastases (*z*) cells. Our central aims are: (i) to demonstrate how decreasing immune surveillance (control parameter I) triggers a defined dynamical route, through limit cycles to chaotic Shilnikov-type bifurcations, that mirrors clinical progression; and (ii) to introduce and validate a quantitative thermodynamic metric, the dissipation function Ψ, derived from a two-factor scheme integrating metabolic flux and dynamical sensitivity, which bridges our dynamical analysis with clinical staging and redefines the systemic robustness of the metastatic state.

The paper is organized as follows: In Sec. 2, we present the network model for gastric cancer metastasis. Section 3 focuses on the mathematical analysis of the model, including stability and bifurcation analysis. Section 4 develops the core thermodynamic framework based on the dissipation function Ψ and its clinical implications. Finally, concluding remarks and an outlook for future research are presented.

## 2. A network model of gastric cancer metastasis

Tumor metastasis is a multi-step process by which tumor cells disseminate from their primary site and form secondary tumors at a distant site. Based on a principle of parsimony and without loss of generality, we proposed a heuristic three-compartment model to capture the essential nonlinear dynamics of gastric cancer metastasis. The system conceptualizes the populations of: primary gastric tumor (*x*), hepatic metastases (*y*), and peritoneal metastases (*z*) cells as interacting populations in a network of biochemical reactions far from thermodynamic equilibrium. Their coupling allows for growth, dissemination, and immune-mediated control, with the immune surveillance strength represented by a key bifurcation (or control) parameter I. The model’s structure is illustrated in the graph shown in Fig. 1.

**Fig. 1.**
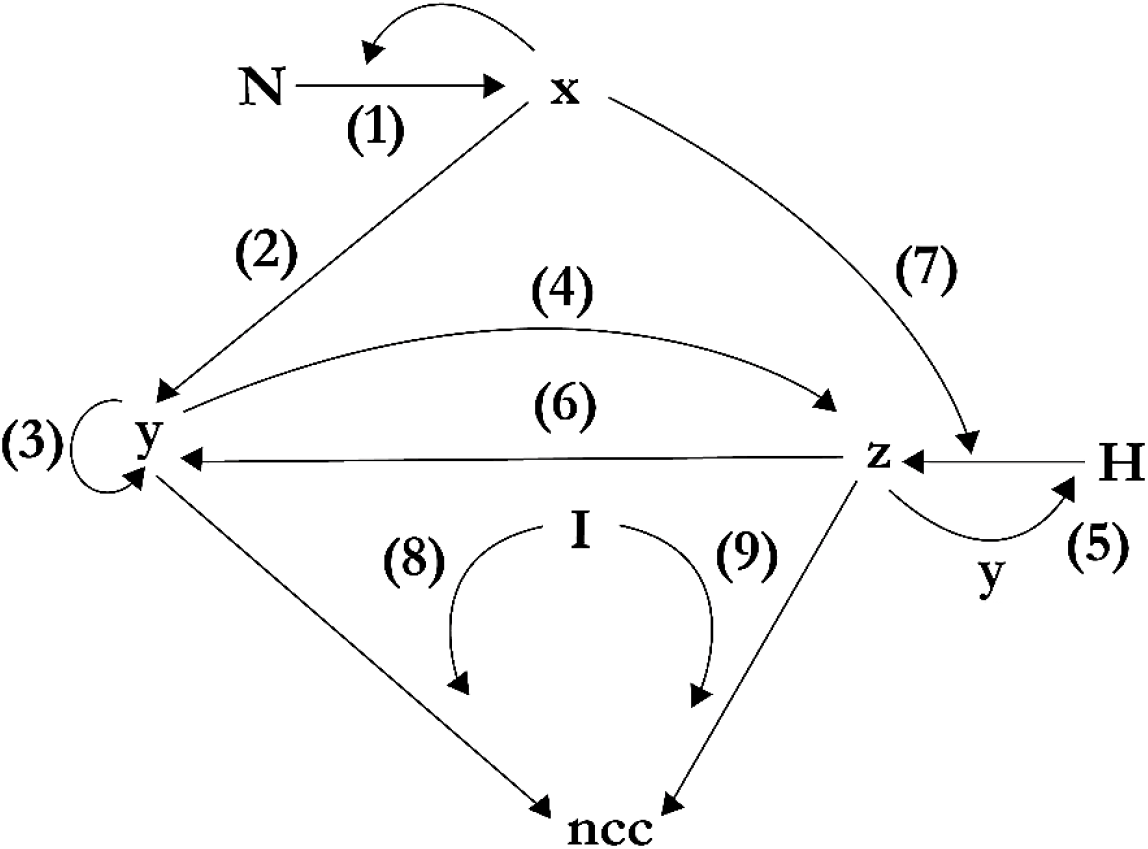
The network model of gastric cancer metastasis

As can be seen, it is based on nine processes that mimic hypothetical nonlinear biochemical reactions and feedback processes. In the model, N represents the population of normal cells exposed to the pro-carcinogenic stimulus; H the population of the host cells in the surrounding environment [35], comprising exclusively epithelial cells; I represents the population of cytotoxic immune cells, specifically cytotoxic T lymphocytes (CTLs) and natural killer (NK) cells, responsible for the adaptive immunosurveillance of metastatic niches [3619], finally, *ncc* represents a population of non-cancerous cells.

The population of primary gastric tumor (*x*) cells can proliferate, seed hepatic metastases (*y*) via hematogenous dissemination, and be eliminated by immune vigilance. The hepatic compartment (*y*) can self-expand and seed the peritoneal compartment (*z*). Conversely, the peritoneal compartment (*z*) can synergistically enhance its own growth and, under a hostile microenvironment H, even repopulate the liver. Critically, both metastatic compartments are subject to adaptive immune control, the efficacy of which is governed by the bifurcation parameter I.

The metastatic cascade is modeled as a network of nine coupled biochemical reactions (see Fig. 1), translating key pathophysiological processes into a dynamical system. The reactions are grouped into three core modules:

### Module A: Primary Tumor Dynamics & Initiation of Dissemination

- *Autonomous Primary Tumor Proliferation*, reaction (1), this reaction captures the uncontrolled, substrate-driven growth of the population of primary gastric tumor (*x*) cells, a fundamental hallmark of cancer [37]. It represents a far-from-equilibrium dissipative process where nutrient consumption fuels exponential biomass increase, modeled by autocatalytic kinetics.
- *Hematogenous Metastatic Dissemination*, reaction (2), this models the intravasation of tumor cells into circulation, a critical step in metastasis. The nonlinear dependence (*x*^2^) represents a threshold or cooperative effect where a critical tumor mass or invasive phenotype is required to trigger systemic spread.

### Module B: Metastatic Niche Expansion and Peritoneal Seeding

- *Hepatic Metastatic Niche Expansion*, reaction (3), this represents the successful colonization and clonal expansion of cancer cells within the liver. It signifies the transition from dormant disseminated cells to actively proliferating metastases, a key event in establishing a robust secondary tumor.
- *Progression to peritoneal carcinomatosis*, reaction (4), this process simulates transcelomic dissemination, a direct anatomical pathway for peritoneal seeding in gastric cancer. The low probability (small *k*_4_) reflects the biological inefficiency of this direct seeding pathway compared to hematogenous dissemination.
- *Microenvironment-Dependent Metastatic Synergy*, reaction (5), this formalizes the “seed-and-soil” hypothesis [38] through an autocatalytic loop. It captures how established metastases (*y, z*) and the host microenvironment H synergistically create a pro-tumor niche that fuels its own expansion, a key driver of aggressive progression.
- *Hostile Microenvironment-Induced Phenotypic Switching*, reaction (6), this reaction models cellular plasticity and adaptive resistance. Under stress from a hostile host environment H, peritoneal cells (*z*) can undergo phenotypic switching, potentially reverting to a state capable of recolonizing other sites (*z*), increasing system resilience.

### Module C: Immune Surveillance and Control

- *Non-Specific Host Response and Immunosurveillance*, reaction (7), this represents the combined, saturable pressure of innate immunity and the host stroma H [35] on the primary tumor. This process involves non-specific killing and physical constraints, a form of biological control distinct from adaptive immunity.
- *Adaptive Immune Control of Metastases*, reactions (8,9), this explicitly models antigen-specific cytotoxicity by T cells and NK cells (control parameter I). It allows the model to account for site-specific differences in immune infiltration and efficacy, crucial for modeling differential responses to immunotherapy.

The constants and concentration of species for the model proposed (see Fig. 1) were chosen empirically, dimensionless, and trying to have as great generality and simplicity as possible, so we have, **Module A:** *k*_1_ = 2, represents the high, substrate-driven efficiency of uncontrolled cancer cell division, the dominant process fueling initial tumor growth; *k*_2_ = 0.5, represent the substantial but less efficient process of intravasation, requiring cells to acquire an invasive phenotype and survive in circulation.

**Module B:** *k*_3_ = 4, it reflects the potential for rapid clonal expansion once disseminated tumor cells successfully colonize a permissive organ niche; *k*_4_ = 0.035, captures the low probability or efficiency of direct transcoelomic seeding from liver metastases or the primary tumor; *k*_5_ = 0.5, characterizes the moderate strength of positive feedback loops in the metastatic niche, where established lesions promote further growth; *k*_6_ = 0.0003, denote the rare event of phenotypic plasticity where peritoneal cells adapt to stress by switching to a liver-colonizing phenotype.

**Module C:** *k*_7_ = 1, sets baseline efficiency for innate immune and stromal pressure. Its impact is scaled by the host factor (H) concentration; *k*_8_ = *k*_9_ = 1, defines the intrinsic efficacy of cytotoxic immune cells (CTL/NK). The overall control strength is modulated by the key bifurcation parameter [I]_CP_, immune activity; [N] = 5 ; [H] = 3 ; [I]_CP_ = [5 −1].

The rate constants *k*_1_ to *k*_9_ were assigned dimensionless values based on their relative biological efficiencies, as informed by the qualitative literature on metastatic cascades [39 Hanahan & Weinberg]. For instance, a high *k*_3_ reflects the rapid clonal expansion in hepatic niches, while a low *k*_6_ models the rarity of phenotypic switching. Crucially, a comprehensive sensitivity analysis [40] was performed across a physiologically plausible hyperparameter space (e.g., varying each *k*_*i*_ by ±50%). This analysis confirms that the qualitative sequence of dynamical transitions, from stable focus to limit cycle to Shilnikov-type chaos with decreasing [I]_CP_, is preserved over a wide parameter range (see Fig. 3.1C). The specific numerical values used thus provide a representative, rather than unique, instantiation of this robust dynamical pathway.

## 3. Mathematical model, stability analysis, and numerical simulations

To translate the conceptual network model (Fig. 1) into a quantitatively analyzable system, we formalize the nine biochemical reactions using the mathematical framework of chemical kinetics. This formalization allows us to derive a system of ordinary differential equations (ODEs) that captures the dynamic interplay between the tumor compartments, their metabolic fluxes, and the critical control exerted by immune surveillance.

Mathematical models represent a suitable way for formalizing the knowledge of living systems obtained through a Systems Biology approach [20,21]. Mathematical modeling of tumor growth makes possible the description of its most important regularities, and it is useful in providing effective guidelines for cancer therapy, drug development, and clinical decision-making [41,42,43].

The model shown in the graph (see Fig. 1) used mathematical methods of chemical kinetics [44] to reduce the network to a system of ordinary differential equations (ODE). The Ordinary Differential Equations (ODE) systems, Eq. (3.1), describe gastric cancer metastasis as:

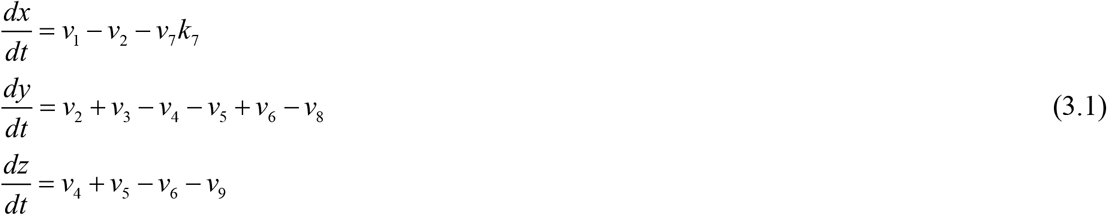

where *v*_*i*_ represent the metabolic fluxes, such as: *v*_1_ = *k* N*x* ; *v*_2_ = 2*k*_2_ *x*^2^ ; *v*_3_ = *k*_3_ *y* ; *v*_4_ = *k*_4_ *y* ; *v*_5_ = *k*_5_ H*yz* ; *v*_6_ = *k*_6_ H*z* ; *v*_7_ = *k*_7_ H*xz* ; *v*_8_ = *k*_8_ I*y* ; *v*_9_ = *k*_9_ I*z*.

Fixed points, stability analysis, bifurcations, spectrum of Lyapunov exponents and sensitivity analysis were calculated using the standard procedure [45,46,47,48,49]. The bifurcation parameter [I]_CP_ represents the population of cytotoxic immune cells, specifically cytotoxic T lymphocytes (CTLs) and natural killer (NK) cells, responsible for the adaptive immunosurveillance of metastatic niches [36,19].

Lyapunov dimension *D*_*L*_, Eq. (2), also known as Kaplan–Yorke dimension [50], was evaluated across the spectrum of Lyapunov exponents *λ*_*j*_ as:

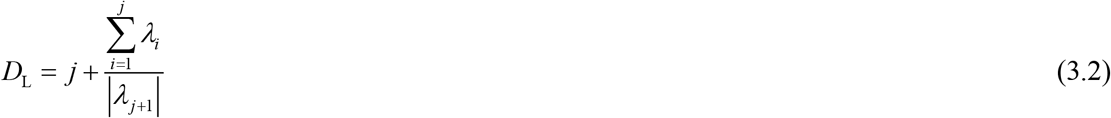

where *j* is the largest integer number for which *λ*_1_ + *λ*_2_ + … + *λ* _*j*_ ≥ 0.

For simulation of the network model, evaluating the spectrum of Lyapunov exponents and sensitivity analysis, COPASI v.4.44.295 software was used [51]. In Table 1, we show the dynamical behavior of the proposed ODEs (3.1) for different values of the control parameter [I]_CP._

**Table 1.** Stability and complexity for the system of ODEs (3.1) for different values of the control parameter [I]_CP_ = [5 −1].

| $[I]_{CP}$ | Eigenvalues of the Jacobian matrix $\varepsilon_i$ | Lyapunov exponents $\lambda_j$ | Lyapunov dimension $D_L$ |
| --- | --- | --- | --- |
| 5<br>ss<br>stable focus | $\varepsilon_{1,2} = -0.077 \pm 5.44i,$<br>$\varepsilon_3 = -8.38$ | $\lambda_1 = -0.0750,$<br>$\lambda_2 = -0.0798,$<br>$\lambda_3 = -8.3760$ | 0 |
| 4.5<br>Limit cycle | $\varepsilon_{1,2} = 0.036 \pm 5.31i,$<br>$\varepsilon_3 = -7.88$ | $\lambda_1 = 0.0,$<br>$\lambda_2 = -0.0706,$<br>$\lambda_3 = -7.6344$ | 1 |
| 1<br>Shilnikov's chaos | $\varepsilon_{1,2} = 0.35 \pm 2.91i,$<br>$\varepsilon_3 = -3.42$ | $\lambda_1 = 0.0401,$<br>$\lambda_2 = 0.0,$<br>$\lambda_3 = -2.3579$ | 2.016 |

In Fig. 2, the dynamic behavior of the network model is shown. It is observed that for low values of the control parameter [I]_CP_ = 1, the tumor cells exhibit “apparently random behavior” (remnant of Shilnikov’s chaos), (see Table 1) [52]. This behavior has important biological implications. On the one hand, the high sensitivity of the system to initial conditions makes long-term predictions unfeasible, i.e., the end forecasts are improbable (poor prognosis). Furthermore, the system displays a high degree of complexity. This implies cancer cells are resilient with respect to pharmacological treatment, thus leading to a low response rate [53].

**Fig. 3_1.**
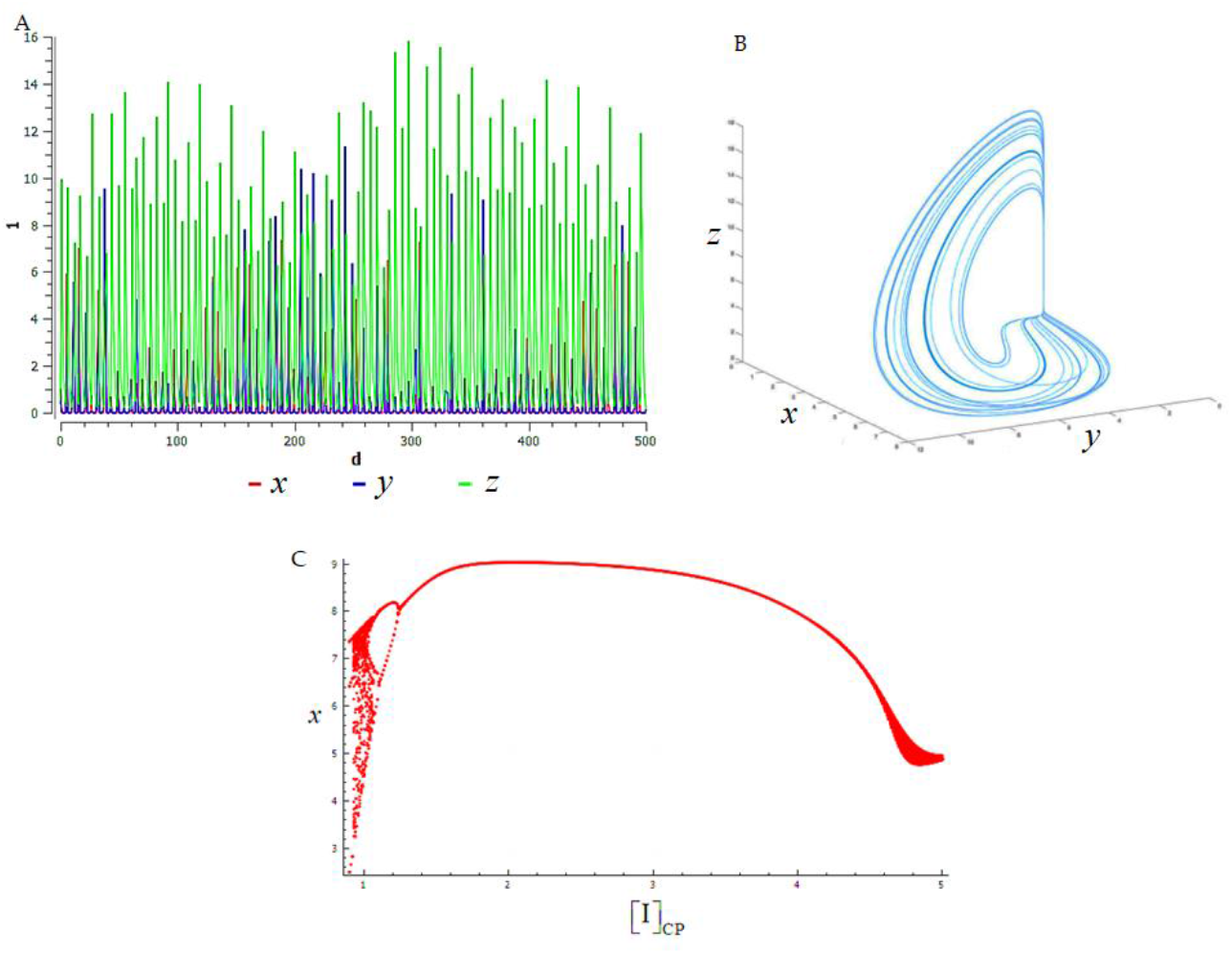
Dynamical behavior of the proposed model (3.1), [I]_CP_ = 1: **A**. time series; **B**. Chaotic attractor; **C**. Bifurcation diagram involves an inverse Feigenbaum (a cascade of saddle-foci Shilnikov’s bifurcations) scenario to evaluate the behavior of the system under a variation of the control parameter [I]_CP._

As can be seen (see Fig. 2C), as immune surveillance declines, take as the control parameter I, produces an inverse Feigenbaum (a cascade of saddle-foci Shilnikov’s bifurcations) scenario, which results in the stabilization of the dynamics and a decrease in the complexity of the system.

Our model suggests that the development of a primary tumor and the subsequent appearance of metastases is not simply the accumulation of malignant cells, but a dynamic, nonlinear, and self-organizing process, far from thermodynamic equilibrium, that exhibits a high degree of robustness, complexity, and hierarchy [20,21] and that, in turn, gives rise to the creation of new information and learning capacity. The information created during the evolutionary process of cancer cannot be destroyed [54], which is manifested by clinical recurrence (relapses) of cancer after a while that has apparently been ‘‘removed’’.

The transition to a chaotic attractor signifies more than complex dynamics; it marks the emergence of a robust, resilient systemic state that underlies treatment resistance and recurrence. This dynamical robustness implies a form of systemic memory or hysteresis, where the state of the system is not easily reversed. In gastric cancer, this is clinically manifested as recurrence (relapse), a fundamental challenge where the disease reappears after apparent remission. A prime example is the high rate of peritoneal recurrence following curative-intent surgery, which remains a leading cause of mortality and is notoriously difficult to treat [55]. Similarly, hepatic recurrence after resection of metastases is common and portends a poor prognosis [56].

This persistence underscores that therapeutic strategies based solely on cytoreduction (mass reduction) are insufficient against a system entrenched in a robust chaotic attractor. A promising complementary approach, supported by studies in other cancers, involves strategies to reprogram the system’s state, altering its dynamics to steer it away from the metastatic attractor [57]. For instance, in gastric cancer, modulating key pathways involved in peritoneal dissemination (such as those governing EMT or mesothelial adhesion) could represent a therapeutic axis informed by this dynamical perspective [58 Nuestro trabajo physA VI y 59 ferroptosis].

Thus, having established the defined route to dynamical chaos driven by declining immune surveillance, a critical question remains: how can one quantify the robustness of this metastatic state? The chaotic attractor itself, while descriptive, does not provide a scalar measure of the system’s thermodynamic resilience. In the following section, we bridge this gap by introducing a coarse-grained dissipation function, Ψ (I), designed to measure the systemic energy dissipation that underlies and sustains this treatment-resistant regime.

## 4. Thermodynamic framework

Non-equilibrium thermodynamics formalism has proven to be both a powerful theoretical framework and a practical tool for understanding and forecasting the emergence and evolution of tumor growth [60,61,62,63,64,65,66,67].

As we know from classic thermodynamics, if the constraints of a system are the temperature *T* and the pressure *p*, then the entropy production can be evaluated using Gibbs’s energy [68], as:

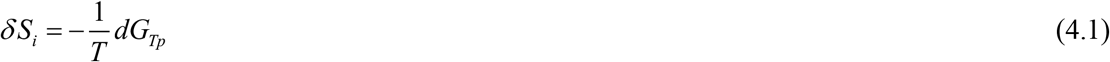

Taking the time derivative of (4.1) yields:

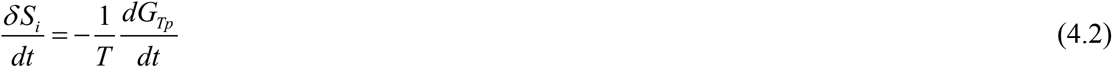

where 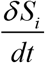 represents the entropy production rate, 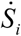. The term 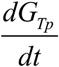 can be developed by means of the chain rule as a function of the degree of advance of the reaction *ξ* as: 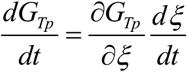. According to De Donder and Van Rysselberghe [69], 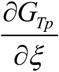 represents the affinity 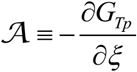, while 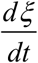 is the reaction rate 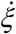. Using the expression for the Gibbs’ equation, *G* = *H* − *TS*, we obtain,

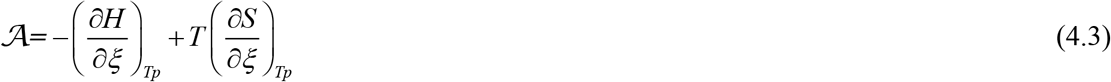

The term 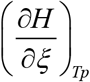 represents the heat of the process, 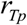. The entropic term 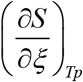 can often be neglected when 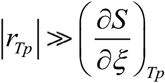 [68]. Considering this, Eq. (4.2) can be rewritten as:

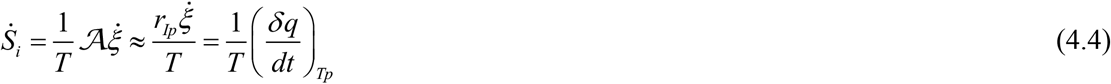

Formula (4.4) provides an approximation for determining the entropy production rate in a living organism [68]. Following Zotin [70], Eq. (4.4) can be expressed in terms of the total metabolic rate 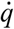 as follows

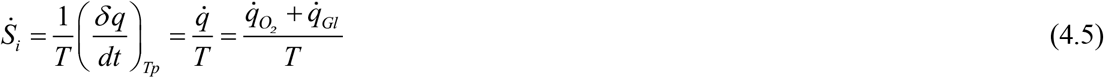

Here, 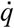 is the total metabolic rate, comprising contributions from oxidative phosphorylation 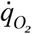 and glycolysis 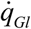. In cancer cells, glycolysis is the dominant process [39]. Consequently, for cancer cells, Eq. (4.5) can be simplified to:

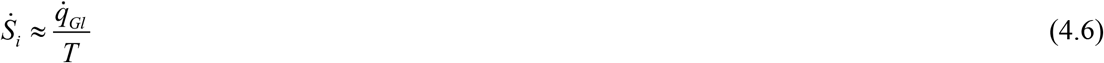

Within the framework of non-equilibrium thermodynamics, it is often convenient to use the dissipation function, 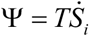, instead of the entropy production rate itself [67]. This function measures the energy per unit time dissipated by a system to maintain its state far from thermodynamic equilibrium and converts the entropy production rate into a non-equilibrium thermodynamic potential. For a complex, open system like metastatic cancer, this function is expected to be high, reflecting the sustained metabolic work required for invasion, adaptation, and immune evasion.

Building on the physical analogy presented in Eq. (4.6), we propose a heuristic *ansatz* to approximate the dissipation function Ψ for our model system. We posit that Ψ (I) can be estimated by a weighted sum of the key metabolic fluxes Φ_*x*_ driving each functional module (**A**:initiation/dissemination, **B**: expansion/synergy, **C**: immune control), normalized by the total metabolic flux:

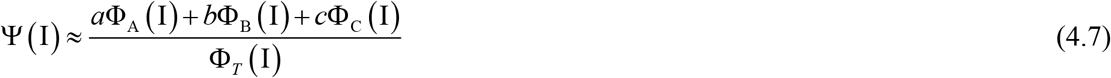

The weighting coefficients *a* (I), *b* (I), *c* (I) in Eq. (4.7) are derived from a two-factor scheme that integrates both the steady-state metabolic activity and the dynamical influence of each functional module, both evaluated as functions of the control parameter I.

First, the module-specific metabolic fluxes Φ _*x*_ (I) are computed at steady state by summing the absolute values of the relevant reaction fluxes *v*_*i*_ listed in Eq. (3.1). For example, Φ_*x*_ (I) = *v*_1_ + *v*_2_, representing the total metabolic throughput of the dissemination module. Second, to account for each module’s dynamical importance, we performed a steady-state sensitivity analysis using COPASI [51]. This yields normalized sensitivity coefficients *s*_*ij*_ (I), quantifying the relative change in each tumor compartment *j* ∈ {*x, y, z*} with respect to perturbations in each rate constant *k*_*i*_. The aggregate dynamical influence of each module is then calculated by summing the absolute sensitivity over its reactions and all compartments. For Module A:

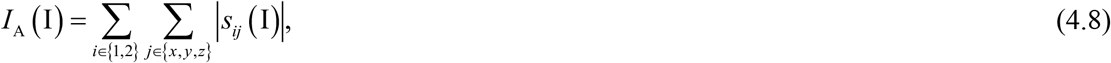

with analogous expressions *I*_B_ (I) and *I*_C_ (I) for Modules B (reactions 3–6) and C (reactions 7–9) respectively.

Finally, the weighting coefficients are obtained by combining these metabolic and dynamical contributions and normalizing:

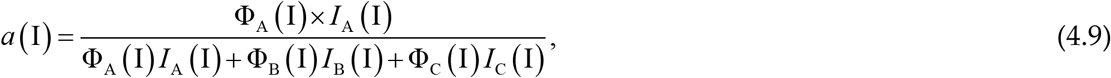

and similarly, for *b* (I) and *c* (I). This hybrid scheme ensures that Ψ (I) reflects not only the raw metabolic output of each module but also its reconfigurable dynamical role in sustaining the metastatic state. Consequently, Ψ (I) quantifies the strategic allocation of metabolic resources to the modules that are most critical for systemic robustness at a given level of immune surveillance I.

In summary, the control parameter I orchestrates a profound reconfiguration of this dynamical landscape. As I decreases, simulating weakened immune surveillance, two interdependent effects emerge: (1) the absolute metabolic fluxes Φ _*x*_ (I) are redistributed, with a relative suppression of immune-mediated control (Module C), and (2) the sensitivity structure is fundamentally altered. The system’s state becomes orders of magnitude more sensitive to perturbations in the core reactions of dissemination (Module A, e.g., *ν*_2_), while the relative influence of modules associated with growth (B) and immune control (C) diminishes. The coefficients *a* (I), *b* (I), *c* (I), *c*are obtained by normalizing the aggregated sensitivity for each module (A: reactions 1-2; B: reactions 3-6; C: reactions 7-9). Therefore, the increasing weight *a* (I) with declining I directly quantifies how the metastatic system’s robustness becomes dynamically dependent on its dissemination capacity, transforming a dynamical stability property into a thermodynamic weighting scheme.

Since all metabolic fluxes Φ_*T*_ (I) share the same units [*d* ^−1^], the proposed dissipation function Ψ is a dimensionless quantity. It does not represent the absolute power dissipation (which would have units of energy per time), but rather the dissipation density or allocation efficiency: the fraction of the total metabolic flux that is strategically allocated, via the weighted coefficients, to maintain the system’s robust, non-equilibrium state. In Table 2, we show the weighting coefficients, metabolic fluxes, and dissipation function Ψ (I) for different values of the control parameter [I]_CP._

**Table 2.** Coefficients, metabolic fluxes, and dissipation function Ψ (I) for different values of the control parameter [I]_CP_ = [5 −1].

| $[I]_{CP}$ | $a(I)$ | $\Phi_A(I)$ | $b(I)$ | $\Phi_B(I)$ | $c(I)$ | $\Phi_C(I)$ | $\Phi_T(I)$ | $\Psi(I)$ |
| --- | --- | --- | --- | --- | --- | --- | --- | --- |
| 5 | 0.35 | 50.5146 | 0.30 | 21.7149 | 0.35 | 49.9829 | 132.2123 | 0.3418 |
| 4.5 | 0.40 | 54.1715 | 0.25 | 20.2062 | 0.35 | 46.3639 | 120.7415 | 0.3558 |
| 1 | 0.50 | 14.3176 | 0.20 | 5.5398 | 0.30 | 15.1653 | 35.0227 | 0.3659 |

As evidenced in Table 2, the coefficients demonstrate a progressive shift in weight toward Module A (dissemination) as I decreases. This reconfiguration aligns with the sensitivity analysis, confirming that in the chaotic metastatic regime, the system becomes highly sensitive to perturbations in dissemination pathways, which form the primary scaffold of its systemic robustness.

The monotonic increase of Ψ (I) (from 0.3418 to 0.3659) as immune surveillance declines confirm our central hypothesis. This trend signifies an enhancement of the system’s dissipative efficiency. The tumor allocates a larger fraction of its diminishing total metabolic flux to sustain the robust (chaotic) dissipative structure characteristic of advanced disease.

The value of Ψ (I) serves as an intensive metric of dissipation density (robustness per unit of metabolic flux). Its increase quantifies a thermodynamic phase transition to a metastatic state that is simultaneously more robust and more refractory to treatment, as its stability becomes less dependent on volumetric growth and more reliant on adaptive dissemination and plasticity.

This numerical progression (0.3418 → 0.3558 → 0.3659) provides a quantitative foundation for correlating thermodynamic robustness with advanced clinical stages (TNM III–IV) [71]. It thereby substantiates the proposed therapeutic paradigm shift: effective intervention in advanced metastasis must pivot from merely reducing tumor mass to strategically destabilizing this robust dissipative structure.

## 5. CONCLUSIONS AND OUTLOOK FOR FUTURE RESEARCH

This work establishes a novel physics-based framework for understanding gastric cancer metastasis by integrating non-equilibrium thermodynamics and nonlinear dynamics. Through a three-compartment heuristic model, we have demonstrated that metastatic progression follows a well-defined dynamical route to chaos. As immune surveillance (parameter I) declines, the system undergoes a supercritical Andronov-Hopf bifurcation, giving rise to limit cycles, followed by a Shilnikov-type saddle-foci bifurcation cascade leading to chaotic attractors. This path mirrors the clinical transition from localized disease to treatment-resistant, systemic metastasis.

In summary, the core theoretical contribution is the introduction of a spatially coarse-grained dissipation function, Ψ (I), constructed via a two-factor ansatz that integrates metabolic fluxes with dynamical sensitivity analysis. This function quantifies the thermodynamic robustness of the metastatic state. We have shown that Ψ (I) increases monotonically as I decreases, validating it as a direct metric of the enhanced dissipative efficiency that characterizes the chaotic, advanced regime. This provides a quantitative bridge between nonlinear dynamical analysis and clinical oncology, showing a decisive correlation with advanced TNM stages (III–IV).

Central to this framework is the dissipation function Ψ (I), a coarse-grained thermodynamic metric derived via a sensitivity-weighted sum of metabolic fluxes. Crucially, Ψ (I) is not a phenomenological construct but is grounded in the system’s reconfigurable dynamical architecture. The monotonic increase of Ψ (I) with declining immune surveillance I validates it as a direct measure of emergent thermodynamic robustness. This increase is mechanistically explained by the sensitivity analysis: as the system transitions to chaos, its stability becomes predominantly governed by the dissemination module (A), a shift captured quantitatively by the recalibrated weighting coefficients *a* (I), *b* (I), *c* (I). Thus, Ψ (I) successfully bridges nonlinear dynamics and clinical staging, providing a physics-based rationale for the therapeutic paradigm shift, from targeting tumor mass to destabilizing the robust dissipative structure it sustains.

This correlation, quantified and mediated by the dissipation function Ψ (I), underscores a fundamental clinical implication: our work advocates for a fundamental paradigm shift in the therapeutic strategy against advanced metastatic cancer. The monotonic increase of Ψ (I) quantifies the emergence of a robust, treatment-resistant dissipative structure. Consequently, effective intervention must pivot from a primarily cytoreductive doctrine, aimed at mass reduction, to a dynamic destabilization strategy. The primary therapeutic objective becomes the reduction of the system’s thermodynamic robustness, i.e., lowering Ψ (I) by strategically perturbing the dominant module, in the metastatic regime, the dissemination network (Module A). This could involve targeting invasive phenotypes, migratory pathways, or the adaptive plasticity that underpins the system’s dissipative efficiency, thereby preventing recurrence by making the metastatic state dynamically untenable.

This dynamical perspective leads to a fundamental reconceptualization of metastatic robustness. The profound adaptability that makes metastasis robust also defines its critical weakness. Each evolutionary step toward spread and colonization, stress resistance, niche dependence, and systemic communication creates a targetable vulnerability. Thus, therapeutic success lies not in attacking bulk tumor growth, but in precisely destabilizing these adaptive supports, turning the metastatic process against itself. Its strength is inherently fragile.

Finally, Outlook for Future Directions:

1. Experimental Validation of Ψ (I) as a Biomarker and Target: Preclinical models are needed to measure metabolic fluxes and perturbation sensitivities *in vivo* to empirically calibrate Ψ (I). This would test whether combined therapies designed to reduce Ψ (I) e.g., anti-dissemination agents (targeting Module A) coupled with microenvironment modulators, can destabilize established metastatic networks and improve treatment efficacy.
2. Model Refinement: Incorporating spatial heterogeneity, stochasticity, and adaptive therapy schedules would enhance the model’s biological realism and clinical translatability.
3. Generalization and Clinical Translation: Applying this approach from the thermodynamics of complex systems to other types of cancer could reveal universal principles of metastatic progression. Ultimately, this contributes to a more fundamental, physics-informed understanding of oncology, where therapeutic strategies are designed not merely to kill cells, but to reconfigure the dysfunctional dissipative dynamics of the tumor-host system.

## ACKNOWLEDGMENTS

To Prof. Dr. Germinal Cocho and Prof. Dr. A. Alzola *in memoriam*. J.M.N-V. is grateful to Alma Mater, Universidad de La Habana.

## DECLARATIONS

### Author Contributions (CRediT)

All authors contributed equally to all aspects of this work, including conceptualization, methodology, software, validation, formal analysis, investigation, resources, data curation, writing, original draft, writing, review & editing, visualization, supervision, and project administration.

### Declaration of Competing Interest

The authors declare that they have no known competing financial interests or personal relationships that could have appeared to influence the work reported in this paper.

### Funding Statement

This research did not receive any specific grant from funding agencies in the public, commercial, or non-profit sectors.

### Declaration of Generative AI and AI-assisted Technologies in the Writing Process

During the preparation of this work, the authors used Grammarly AI for language polishing and editorial assistance. After using this tool, the authors reviewed and edited the content as needed and take full responsibility for the content of the published article.

## Notes

### Competing Interest Statement

The authors have declared no competing interest.

